# Alkaline phosphatase activity of serum affects osteogenic differentiation cultures

**DOI:** 10.1101/2021.10.01.462693

**Authors:** Sana Ansari, Keita Ito, Sandra Hofmann

## Abstract

Fetal bovine serum (FBS) is a widely used supplement in cell culture medium, despite its known variability in composition which greatly affects cellular function and consequently the outcome of studies. In bone tissue engineering, the deposited mineralized matrix is one of the main outcome parameters, but using different brands of FBS can result in large variations. Alkaline phosphatase (ALP) is present in FBS. Not only is ALP used to judge the osteogenic differentiation of bone cells, it may affect deposition of mineralized matrix. The present study focused on the enzymatic activity of ALP in FBS of different suppliers and its contribution to mineralization in osteogenic differentiation cultures. It was hypothesized that culturing cells in a medium with high intrinsic ALP activity of FBS will lead to higher mineral deposition compared to media with lower ALP activity. The used FBS types were shown to have significant differences in enzymatic ALP activity. Our results indicate that the ALP activity of the medium not only affected the deposited mineralized matrix but also the osteogenic differentiation of cells as measured by a changed cellular ALP activity of human bone marrow derived mesenchymal stromal cells (hBMSC). In media with low inherent ALP activity, the cellular ALP activity was increased and played the major role in the mineralization process; while, in media with high intrinsic ALP activity contribution from the serum, less cellular ALP activity was measured and the ALP activity of the medium also contributed to mineral formation substantially. Our results highlight the diverse effects of ALP activity intrinsic to FBS on osteogenic differentiation and matrix mineralization and how FBS can determine the experimental outcomes, in particular for studies investigating matrix mineralization. Once again, the need to replace FBS with more controlled and known additives is highlighted.

**Graphical abstract:** 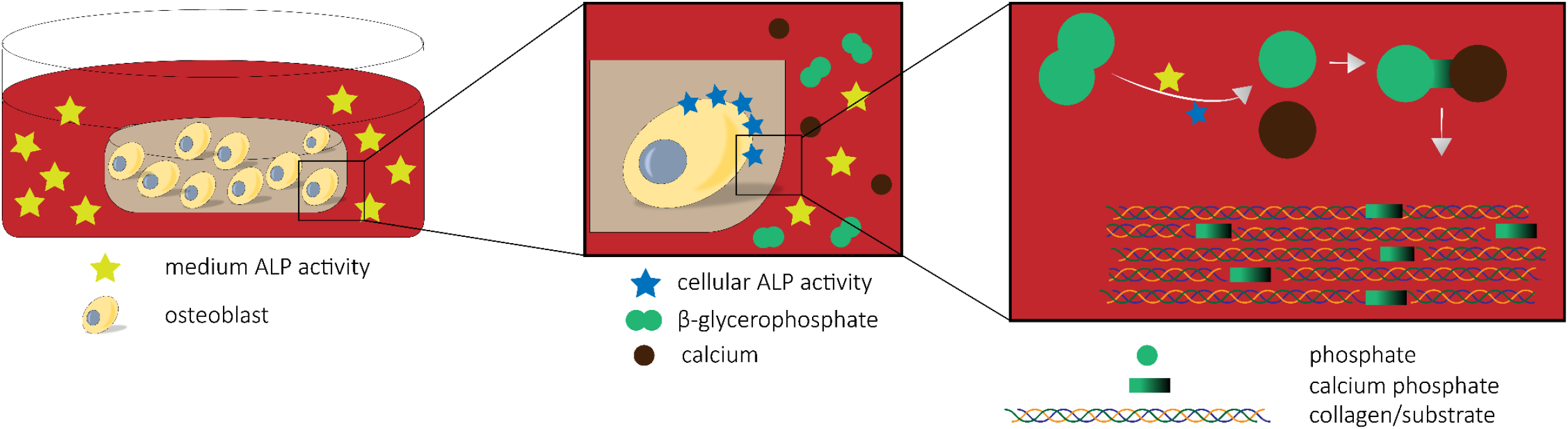

## Introduction

Fetal bovine serum (FBS) is a widely known supplement in cell culture media, used at concentrations up to 20% (v/v) [1]. FBS provides cells with vital factors including growth factors, hormones, and vitamins essential for cell survival, growth, and division [1,2]. However, the use of FBS in *in vitro* cell culture is controversial due to a number of reasons, including ethical concerns, shortage in global supply and most importantly its undefined, complex composition and variability which could lead to unexpected and/or unreliable experimental outcomes [1,3,4]. Thus, either complete avoidance of FBS or at least awareness of the effects that some components of FBS might have on experimental outcomes should be considered [2,5,6].

FBS has previously been described having various effects on mineral deposition. It was shown being able to hydrolyze phosphate sources and by that increasing the concentration of free phosphate in the culture medium, which further resulted in mineralization of fibrous proteins such as collagen and silk fibroin even without the presence of cells [7,8]. Moreover, the deposited calcium content on the fibrous scaffolds was significantly affected by the variation in the chemical composition of FBS [8]. Since the exact chemical composition of FBS is not provided and is known to differ even between batches within the same brand, it remains unknown which component(s) contributes to the mineralization process. On the other hand, knowledge on which and how FBS component(s) contribute to mineralization could be beneficial for *in vitro* studies where mineralization of extracellular matrix is needed (*e*.*g*. bone tissue engineering) but also where mineralization should be avoided (*e*.*g*. cardiac tissue engineering).

Alkaline phosphatase (ALP) is a potential component of FBS affecting mineralization. ALP is an abundant membrane-bound glycoprotein [9]. It exists as four isozymes, depending on the tissue where it is expressed: placental ALP, germ cell ALP, intestinal ALP and liver/bone/kidney ALP [10]. ALP enzymes expressed in the placenta, germinal and intestine tissue are tissue-specific meaning that under physiological conditions, they are found exclusively in the tissues where they are expressed, whereas the ones expressed in liver, bone, and kidney are known as tissue-nonspecific ALP because they can also be found in blood circulation [11–14].

In bone, ALP is expressed by osteoblasts, the bone forming cells, and either anchored to the cell membrane or to matrix vesicles generated by osteoblasts through glycosylphosphatidylinositol (GPI) linkage attached to the carboxyl-terminal of the enzyme [15]. ALP can be released into serum through matrix vesicles or after its cleavage from the osteoblast surface by circulating GPI-specific phospholipase D [14,16,17]. Thus, serum contains ALP which is used for example as a biomarker in the clinics to assess chronic kidney diseases or bone disorders [17].

During the osteogenic differentiation process, the presence and activity of ALP indicates the differentiation of mesenchymal stromal cells (MSCs) towards osteoblasts [18]. The activity of ALP can be measured thorough colorimetric assays where p-nitrophenyl phosphate, a phosphate substrate, is dephosphorylated by ALP [19,20]. Besides the activity of ALP, the expression of ALP can be measured through techniques such as quantitative reverse transcription-polymerase chain reaction (RT-PCR), western blot, and immunofluorescence imaging [21,22]. The latter can determine the location of expressed ALP with respect to the cell.

ALP expressed by osteoblasts is an important enzyme in the process of bio-mineralization [12]. This enzyme can hydrolyze extracellular inorganic pyrophosphate, generated by the hydrolysis of adenosine triphosphate (ATP), which leads to an increase in the local concentration of inorganic phosphate (Pi) [23–26]. Pi and calcium ions are thought to accumulate inside matrix vesicles to form amorphous calcium phosphate or hydroxyapatite crystals which is believed to be the initial stage of extracellular matrix mineralization during bone formation [27].

In *in vitro* bone studies, to avoid spontaneous mineralization, β-glycerophosphate (β-GP) has been used as the phosphate source that is believed to be cleaved through the ALP activity of osteoblasts, making Pi available for matrix mineralization [28]. However, hydrolyzing β-GP under cell-free conditions and in the presence of FBS indicated that serum ALP activity has its contribution in making Pi available in culture medium for subsequent calcium phosphate deposition [7,29]. This effect resulted in non-physiological and uncontrollable mineralization prior to osteoblast differentiation *in vitro* which needs to be avoided in many research lines, for instance, the development of *in vitro* bone models [30].

In this study, four different types of FBS with different intrinsic ALP activity were investigated with the aim to investigate the influence and contribution of medium (provided by FBS) and cellular ALP activity on mineralized tissue formation. For this, silk fibroin scaffolds were either left acellular or were seeded with human bone marrow derived mesenchymal stromal cells (hBMSCs) and cultivated *in vitro*. We hypothesized that the ALP activity of medium containing FBS not only affects calcium phosphate deposition in the presence and absence of cells, but that it also has an influence on the cellular ALP activity. We further investigated whether heat inactivation of FBS, a process which is commonly used to destroy complement activity in serum, also can eradicate the effects of FBS ALP. Knowledge on the influence of ALP activity of FBS, as one of the many components in FBS that could be responsible for the high variation in experimental outcomes, can shed a light on the necessity of developing serum-free medium with clearly defined components.

## Materials and Methods

### 1 Materials

Dulbecco’s modified eagle medium (DMEM high glucose, Cat. No. 41966 and low glucose Cat. No. 22320), antibiotic/antimycotic (Anti-Anti, Cat. No. 15240062), non-essential amino acids (NEAA, Cat. No. 11140050), and trypsin-EDTA (0.5%, Cat. No. 2530054) were from Life Technologies (The Netherlands). FBS types were from Bovogen (Cat. No. SFBS), Sigma (Cat. No. F7524), Hyclone (South American research grade FBS, Cat. No. SV30160.02), and U.S. Origin FetalClone III serum (Fetalclone III, Cat. No. SH30109.03). Silkworm cocoons were purchased from Tajima Shoji Co., LTD. (Japan). Unless noted otherwise, all other substances were of analytical or pharmaceutical grade and obtained from Sigma Aldrich (The Netherlands).

### 2 Measurement of ALP activity of serum and medium supplemented with FBS

The ALP activity of four types of FBS and the resulting control medium containing DMEM low glucose, 1% Anti-Anti and 10% FBS (Table 1) was measured as follows: In a 96-well plate, 80 μL of each serum sample or medium sample were mixed with 20 μL of 0.75 M 2-amino-2-methyl-1-propanol buffer and 100 μL 10 mM p-nitrophenylphosphate solution and incubated until colour developed, before 0.2 M NaOH were added to stop the conversion of p-nitrophenylphosphate to p-nitrophenol. Absorbance was measured in a spectrophotometer at 450 nm and ALP activity was calculated by comparison to standards of known p-nitrophenol concentration.

**Table 1.** List of abbreviations of FBS types and the medium containing each type of FBS

| Serum brand | LOT number | Abbreviation | Medium | Abbreviation |
| --- | --- | --- | --- | --- |
| Bovogen | 51113 | **B** | Control medium containing 10%<br>Bovogen | **DMEM%10B** |
| Sigma | 7611 | **S** | Control medium containing 10%<br>Sigma | **DMEM%10S** |
| Hyclone | RE00000004 | **H** | Control medium containing 10%<br>Hyclone | **DMEM%10H** |
| Fetalclone III | AD19958305 | **F** | Control medium containing 10%<br>Fetalclone III | **DMEM%10F** |

### 3 Heat inactivation of FBS

5 mL of each serum type was placed in a water bath at 56°C for 30 minutes. After 30 minutes, the sera samples were removed from the water bath and transferred into an ice bath for rapid cooling. The ALP activity of heat inactivated (HI) FBS and the media containing 10% of HI FBS was measured according to section 2.

### 4 Measurement of Pi concentration in medium supplemented with FBS

Concentration measurements of free phosphate Pi in control medium containing DMEM low glucose, 1% Anti-Anti and 10% FBS or HI FBS (Table 1 and 2) with and without the addition of 10 mM β-glycerophosphate (β-GP) after 48 hours of incubation at 37°C were performed according to the manufacturer’s instruction (Malachite Green Phosphate Assay Kit, Sigma-Aldrich, The Netherlands). Briefly, 80 μL of 1:200 (v/v) diluted samples in ultrapure water (UPW) were mixed with 20 μL of working reagent and incubated for 30 minutes at room temperature. In this assay, a green complex is formed between molybdate and Pi. Colour formation from the reaction was measured spectrophotometrically at 620 nm and phosphate concentration was calculated by comparison to a phosphate standard provided in the kit.

**Table 2.** List of abbreviations of heat inactivated (HI) FBS and the medium containing each type of HI FBS

| **Serum** | **Abbreviation** | **Medium** | **Abbreviation** |
| --- | --- | --- | --- |
| HI Bovogen | **HI-B** | Control medium containing 10% HI Bovogen | **DMEM%10HI-B** |
| HI Sigma | **HI-S** | Control medium containing 10% HI Sigma | **DMEM%10HI-S** |
| HI Hyclone | **HI-H** | Control medium containing 10% HI Hyclone | **DMEM%10HI-H** |
| HI Fetalclone III | **HI-F** | Control medium containing 10% HI Fetalclone III | **DMEM%10HI-F** |

### 5 Scaffold fabrication

To prepare silk fibroin scaffolds, 3.2 grams of cut and cleaned Bombyx mori L. silkworm cocoons were degummed by boiling in 1.5 L UPW containing 0.02 M Na_2_CO_3_ for 1 hour, whereafter it was rinsed with 10 L cold UPW to extract sericin. Dried purified silk fibroin was dissolved in 9 M lithium bromide (LiBr) solution in UPW at 55°C for 1 hour and dialyzed against UPW for 36 hours using SnakeSkin Dialysis tubing (molecular weight cutoff: 3.5 kDa, Thermo Fisher Scientific, The Netherlands). The silk fibroin solution was frozen at -80°C for at least 2 hours and lyophilized (Freezone 2.5, Labconco, USA) for 4 days. 1.7 grams of lyophilized silk fibroin was then dissolved in 10 mL 1,1,1,3,3,3-Hexafluoro-2-propanol (HFIP) at room temperature for 5 hours resulting in a 17% (w/v) solution. 1 mL of silk-HFIP solution was added to a Teflon container containing 2.5 grams NaCl with a granule size between 250-300 μm. After 3 hours, HFIP was allowed to evaporate for 4 days. Silk fibroin-NaCl blocks were immersed in 90% (v/v) methanol (Merck, The Netherlands) in UPW for 30 minutes to induce the protein conformational transition to β-sheets [31]. Scaffolds were cut into disks of 3 mm height with an Accutom-5 (Struer, Type 04946133, Ser.No. 4945193), followed by immersion in UPW for 2 days to extract NaCl. Disc-shaped scaffolds were made with a 5 mm diameter biopsy punch (KAI medical, Japan) and autoclaved in phosphate buffered saline (PBS) at 121°C for 20 minutes.

### 6 Cellular and acellular scaffolds preparation

Human bone marrow mesenchymal stromal cells (hBMSCs) were isolated from human bone marrow (Lonza, USA) and characterized as previously described [32]. Passage 3 hBMSCs were expanded in expansion medium (DMEM high glucose with 10% FBS Sigma, 1% Anti-Anti, 1% NEAA, and 1 ng/ml bFGF) for 7 days. At day 7, cells were 80% confluent and trypsinized. 16 scaffolds were dynamically seeded with 1*10^6^ cells per scaffold as previously described [33]. Briefly, each scaffold was incubated with a cell suspension (1*10^6^ cells/4 mL control medium (DMEM, 10% FBS respective of each group, 1% Anti-Anti)) in 50 mL tubes placed on an orbital shaker at 150 rpm for 6 hours in an incubator at 37°C [33]. The remaining scaffolds were left acellular and incubated in the control medium as described above. All scaffolds were incubated in 24-well plates at 37°C and 5% CO_2_ for 4 weeks. Each well was filled with 1 mL osteogenic medium (control medium from Table 1 supplemented with 50 μg/mL ascorbic-acid-2-phosphate, 100 nM dexamethasone, 10 mM β-GP). The medium was refreshed 3 days a week.

### 7 Measurement of ALP activity of cells

After 4 weeks of culture, scaffolds (n=3 per group) were washed with PBS and each disintegrated in 500 μL of 0.2% (v/v) Triton X-100 and 5 mM MgCl_2_ solution using steel balls and a minibeadbeater™ (Biospec, USA). The solids were separated by centrifugation (3000 g, 10 minutes). The measurement of ALP activity in the supernatant was performed as described in section 3. In a 96-well plate, 80 μL of the supernatant was mixed with 20 μL of 0.75 M 2-amino-2-methyl-1-propanol buffer and 100 μL 10 mM p-nitrophenylphosphate solution and incubated until colour developed, before 0.2 M NaOH were added to stop the conversion of p-nitrophenylphosphate to p-nitrophenol. Absorbance was measured spectrophotometrically at 450 nm and ALP activity was calculated by comparison to standards of known p-nitrophenol concentration.

### 8 Measurement of (soluble) calcium concentration in medium supplemented with FBS

The calcium concentration was performed on control medium and osteogenic medium in the presence and absence of cells after 48 hours of incubation at 37°C. 5 μL of each medium condition was mixed with 95 μL of calcium working solution (Stanbio Calcium (CPC) LiquiColor® Test, Stanbio Laboratories) and incubated at room temperature for at least 1 minute. In this assay, the calcium ion concentration is measured by the chromogenic complex formed between calcium ions and o-cresolphthalein. Absorbance at 550 nm was measured and calcium concentration was calculated by comparison to standards of known calcium chloride concentrations.

### 9 Measurement of (deposited/precipitated) calcium and phosphate on cell-seeded and acellular scaffolds

After 4 weeks of culture, scaffolds (n=3 per group) were washed with PBS and each disintegrated in 500 μL of 5% Trichloroacetic acid (TCA) in UPW using steel balls and a minibeadbeater™ (Biospec, USA). After 48 hours of incubation at room temperature, the solids were separated by centrifugation (3000 g, 10 minutes). Calcium and phosphate assay were performed on each sample as described below.

#### 9-1 Measurement of (deposited/precipitated) calcium on scaffolds

5 μL of samples were mixed with 95 μL of calcium working solution (Stanbio Calcium (CPC) LiquiColor® Test, Stanbio Laboratories) and incubated at room temperature for at least 1 minute. In this assay, the calcium ion concentration is measured by the chromogenic complex formed between calcium ions and o-cresolphthalein. Absorbance at 550 nm was measured and calcium concentration was calculated by comparison to standards of known calcium chloride concentrations.

#### 9-2 Measurement of (deposited/precipitated) phosphate on scaffolds

A phosphate assay was performed according to the manufacturer’s instruction (Malachite Green Phosphate Assay Kit, Sigma-Aldrich, The Netherlands). Briefly, 80 μL of 1:200 (v/v) diluted samples in UPW were mixed with 20 μL of working reagent and incubated at room temperature for 30 minutes. Absorbance was measured spectrophotometrically at 620 nm and phosphate concentration was calculated by comparison to the phosphate standard provided in the kit.

### 10 Histology

After 4 weeks of culture, scaffolds were washed with PBS and immersed first in 5% and then in 35% sucrose solution in PBS at room temperature for 10 minutes each. The scaffolds were embedded in cryomold containing Tissue-Tek OCT compound (Sakura, The Netherlands), frozen on dry ice, cut into 5 μm thick sections using a cryostat cryotome (Fisher Scientific, The Netherlands) and mounted on Superfrost Plus™ microscope slides (Thermo Fisher Scientific, The Netherlands). Sections were washed with PBS, fixed in 10% neutral buffered formalin for 10 minutes at room temperature, washed again with PBS and stained with Alizarin Red to identify mineralization.

### 11 Micro-computed tomography imaging (μCT)

μCT measurements were executed on a μCT100 imaging system (Scanco Medical, Brüttisellen Switzerland) after 4 weeks of culture (n=4 per group). Scanning was performed at an isotropic nominal resolution of 17.2 μm, an energy level of 55 kVp, and an intensity of 200 μA. Integration time was set to 300 ms and two-fold frame averaging was performed. To reduce part of the noise, a constrained Gaussian filter was applied. Filter support was set to 1.0 and filter width sigma to 0.8 voxel. To distinguish mineralized tissue from non-mineralized tissue, segmentation was performed. A global threshold range was set to 148-1970 after visual judgment of the grey images to identify mineralized structures compared to histologically stained samples. Unconnected objects smaller than 50 voxels were removed through component labelling and neglected for further analysis. Quantitative morphometry was performed to assess the mineralized volume of the entire construct [34].

### 12 Statistics

GraphPad Prism 9.0.2 (GraphPad Software, USA) was used to perform statistical analysis and to make graphs. For Figure 1 A-B and Figure 6 A-B, a Kruskal-Wallis test with Dunn post-hoc testing was performed. Figures 1 C, 2 B, 3, 4 I and 6 C were analysed by Mann-Whitney test. Differences between groups were considered statistically significant at a level of p < 0.05. Histological figures show representative images per group of all the samples assessed.

**Figure 1.**
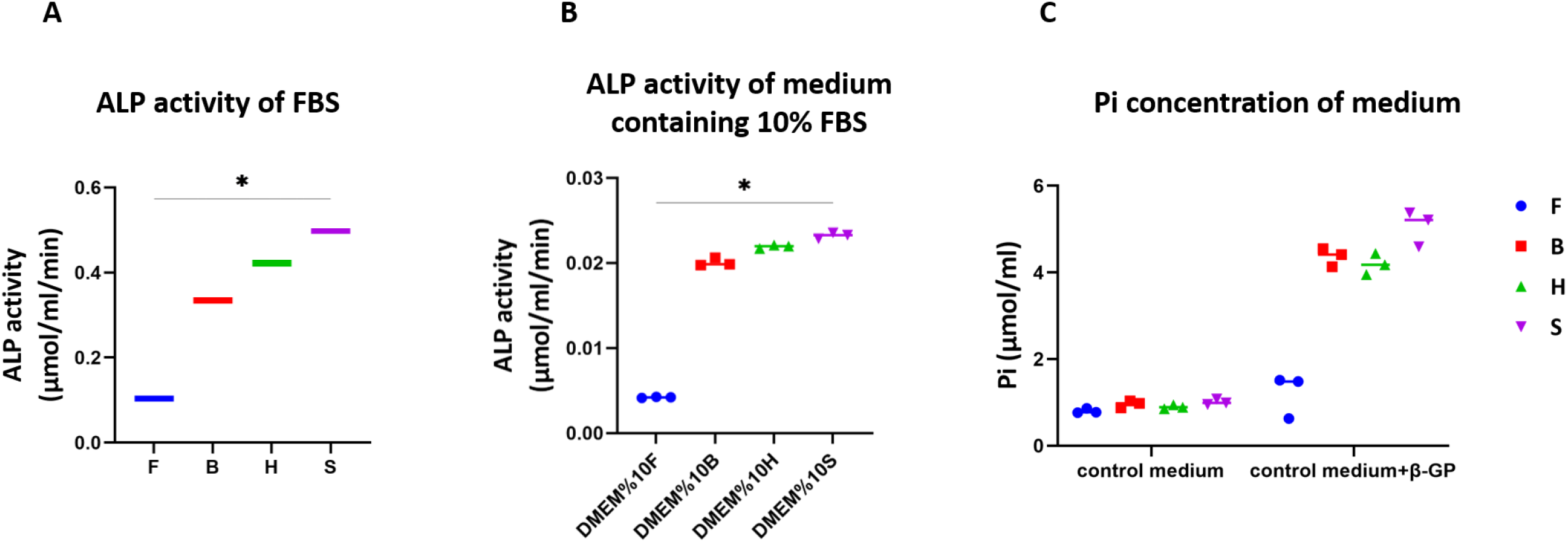
ALP is present in FBS and its activity was different between the four different FBS types tested (A). Control medium supplemented with 10% FBS showed differences in ALP activity with the same trend (B). 48 hours incubation of control medium supplemented with FBS and 10 mM β-GP resulted in an increase in Pi concentration in the medium (C). The increase in Pi seems to be correlated with the ALP activity in the medium, it showed that the FBS with lowest ALP activity (Fetal clone III) led to lowest increase in Pi concentration of the medium.

**Figure 2.**
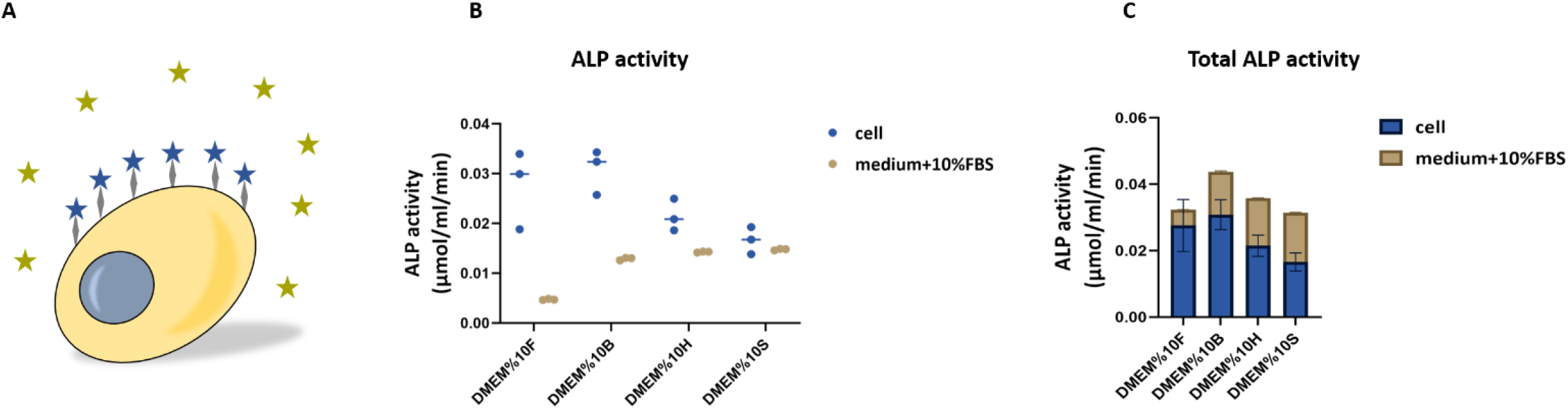
Osteoblasts express ALP, a membrane-bound protein (A-blue stars) and the medium containing FBS has shown to have active ALP (A-dark yellow stars). The cellular ALP activity seems negatively correlated to the medium ALP activity; in the groups with low medium ALP activity, the cells expressed higher ALP activity compared to the groups with high ALP activity (B). The total ALP activity in all groups was equal with no significant differences (C).

**Figure 3.**
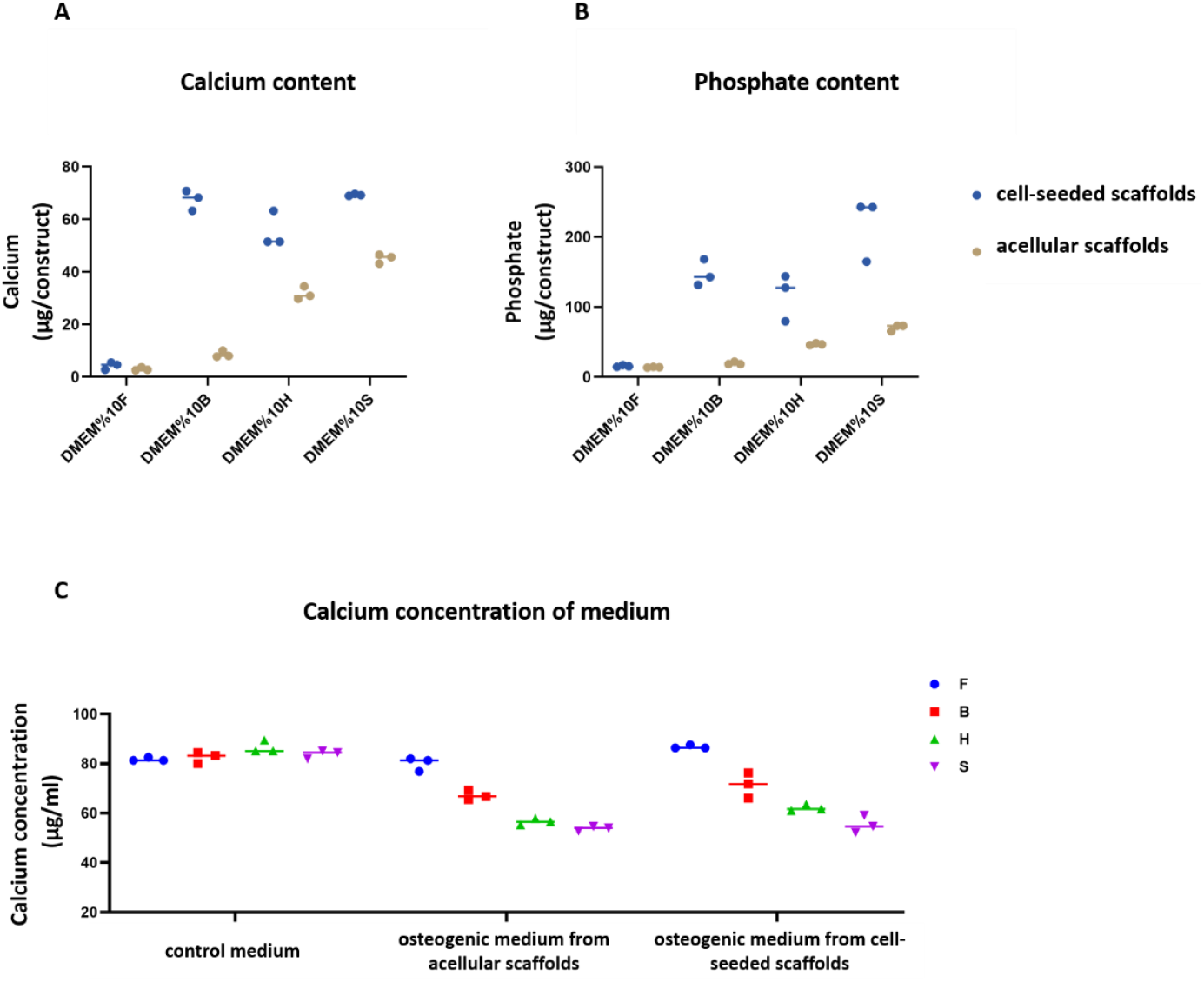
Deposited calcium (A) and phosphate (B) after 4 weeks of culture either without (contribution of medium ALP activity, dark yellow dots) or with cells (contribution of cellular ALP activity, blue dots) on 3D silk fibroin scaffolds. The calcium concentration in the medium was decreased at higher ALP activities, probably because it was deposited in the form of calcium phosphate (C).

**Figure 4.**
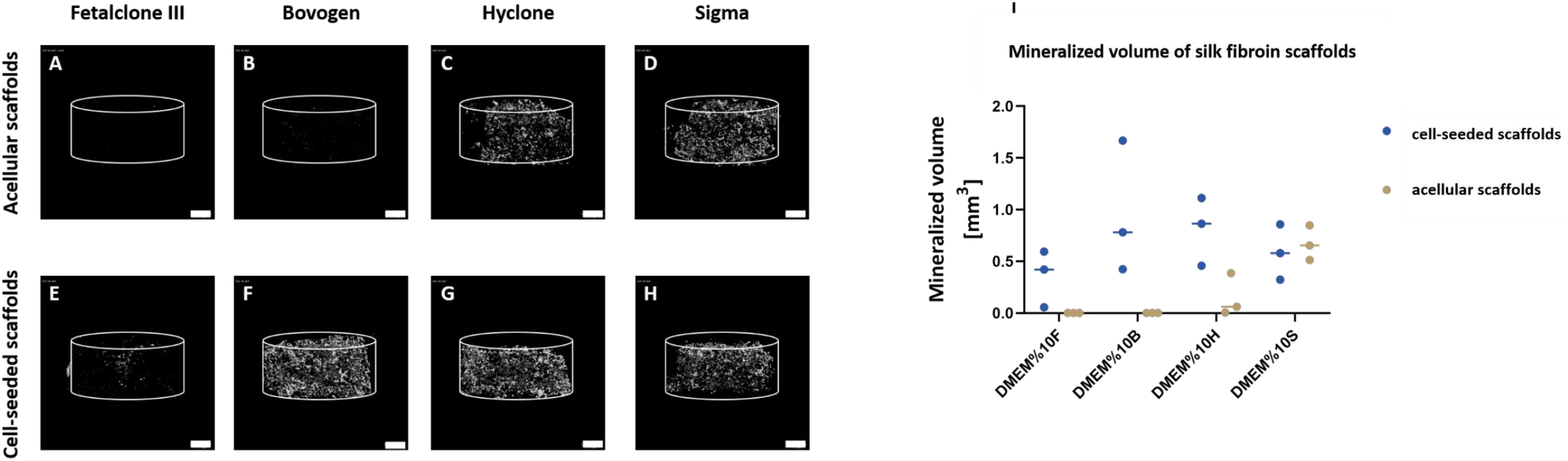
μCT analysis of mineralized volume within acellular and cell-seeded silk fibroin scaffolds after 4 weeks of culture. In the medium containing Fetalclone III (A and E) and Bovogen (B and F) with low medium ALP activity, the mineral deposition happened exclusively in the presence of cells. In the Hyclone and Sigma containing medium groups with high medium ALP activity, substantial amounts of mineral deposition happened even if cells were not present (C, G, D, and H). Scale bar: 1mm.

## Results

### 1 The ALP activity of FBS elevated the Pi concentration in the medium

Four different FBS types and their corresponding control media containing 10% FBS were analysed for their intrinsic ALP activity. The ALP activity between the different brands were different from one another in both concentrated (Figure 1 A) and diluted state (Figure 1 B). As the enzymatic activity is influenced by enzyme concentration, high ALP activity corresponds to high concentration of ALP in FBS [35]. The ALP activity in the diluted state decreased significantly compared to concentrated FBS which was not necessarily 10 times less. To investigate whether this enzymatic activity contributes to the supply of Pi in the medium, control media containing 10% FBS were supplemented with 10 mM β-GP. β-GP is generally used as the phosphate source for *in vitro* osteogenic differentiation processes. The enzymatic activity of ALP was able to convert β-GP into Pi, resulting in an increased Pi level in the medium. The concentration of Pi in the medium was elevated by a factor of 1.51, 4.54, 4.69, and 5.03 fold in medium supplemented with 10% Fetalclone III, Bovogen, Hyclone, and Sigma FBS, respectively, within 48 hours of incubation compared to respective control medium (Figure 1 C). This indicates that the activity of ALP present in FBS is capable to cleave β-GP regardless of the presence of cells in the system. Moreover, the increase in the concentration of Pi after 48 hours was correlated to the intrinsic ALP activity of medium containing FBS.

### 2 Cellular and medium ALP activity was negatively correlated

Two types of ALP activity were measured after 4 weeks of culture since they can both contribute to the overall amount of available Pi. First, the activity of membrane-bound ALP, expressed by osteoblasts during osteogenic differentiation (Figure 2 A, blue stars). Second, the ALP activity present within the different media containing 10% FBS (Figure 2 A, dark yellow stars). The measured ALP activity was normalized to the time of incubation. The cellular enzymatic activity of ALP in the groups of medium containing Fetalclone III and Bovogen was higher than that of cells grown in media containing Hyclone and Sigma FBS. This was in contrast to the activity of ALP in medium containing FBS. There seemed to be a negative correlation between the cellular and medium ALP activity; in media with low inherent ALP activity, the cells have a higher ALP activity (Fetalclone III) compared to the medium with high inherent ALP activity (Sigma) (Figure 2B). However, this was not proportional to the ALP activity of the media. Notably, after four weeks, the total ALP activity in the construct was roughly equal in all four groups and did not show any significant differences (Figure 2C).

### 3 Cellular and medium ALP activity both contribute to calcium phosphate deposition

The amount of calcium and phosphate deposited within the constructs were measured both in the presence and absence of cells after 4 weeks. Incubation of acellular scaffolds in medium containing FBS indicated the contribution of medium ALP activity on calcium phosphate deposition. The ALP activity inherent to the media enabled the deposition of calcium phosphate even when no cells were present. As expected, the amount of calcium and phosphate per construct varied in different culture media used (Figure 4). The deposited calcium phosphate per acellular scaffold (Figure 4 A and B, dark yellow dots) which indicates the contribution of medium ALP activity showed the same trend as the ALP activity of the medium (Figure 3, dark yellow dots): Sigma > Hyclone > Bovogen > Fetalclone III FBS. As hypothesized, even in the absence of cells, a high ALP activity in medium resulted in more calcium phosphate deposition compared to the medium with low ALP activity.

The presence of cells and their differentiation towards osteoblasts increased the calcium phosphate deposition further, which indicated the contribution of cellular ALP activity (Figure 3, blue dots) next to the medium ALP activity. The cellular ALP activity resulted in increasing Pi and thus calcium phosphate deposition. The calcium and phosphate content of cell-seeded scaffolds followed the following pattern: Sigma > Bovogen > Hyclone > Fetalcone III FBS. This pattern is not, however, consistent with the cellular ALP activity which was Bovogen > Fetalclone III > Hyclone > Sigma FBS.

The calcium concentration of control medium in all groups were similar, as expected. When the medium was supplemented with osteogenic factors including β-GP in the absence and presence of cells, this concentration decreased in the medium (Figure 3 C). The medium with high ALP activity (Sigma) showed a larger decrease of calcium concentration in the medium compared to medium with low ALP activity (Fetalclone III). The decrease of calcium concentration in the medium was indicative of the deposited calcium phosphate on the scaffolds.

### 4 μCT analysis and Alizarin red staining detected calcium phosphate deposition on both acellular and cell-seeded scaffolds

μCT imaging (Figure 4) and histology (Figure 5) of the samples after 4 weeks of culture confirmed the deposition of a mineralized matrix either within the silk fibroin scaffold and/or in the extracellular space. Incubation of acellular scaffolds in media containing Hyclone and Sigma with high medium ALP activity led to mineral deposition within the silk fibroin scaffold. The resulting mineral volume was significantly higher than that on scaffolds incubated in media containing Fetalclone III and Bovogen FBS with low medium ALP activity. In the presence of cells, the mineralized volume changed significantly in all groups compared to acellular constructs, most likely as a result of the cellular ALP activity. With μCT, the mineralized volume in the medium containing Fetalclone III and Bovogen FBS was visible only in the presence of cells; while the media containing Hyclone and Sigma FBS showed large mineralized volumes even on acellular scaffolds which indicates the contribution of their medium ALP activity (Figure 4 I). As the scaffolds are made of silk fibroin, which is a protein similar to collagen, in the acellular groups, the minerals are expected to be found in and/or on the scaffolds. In the cell-seeded groups, the minerals could be found both in/on the scaffolds and in the extracellular matrix (ECM) formed by cells. On acellular scaffolds cultured in the medium containing Hyclone and Sigma FBS, minerals were detected on the scaffolds, most likely showing mineral precipitations (Figure 5 C and D). The cell-seeded scaffolds cultured in medium containing Bovogen, Hyclone and Sigma FBS showed mineralization both in the ECM and within the scaffolds (Figure 5 F, G and H).

**Figure 5.**
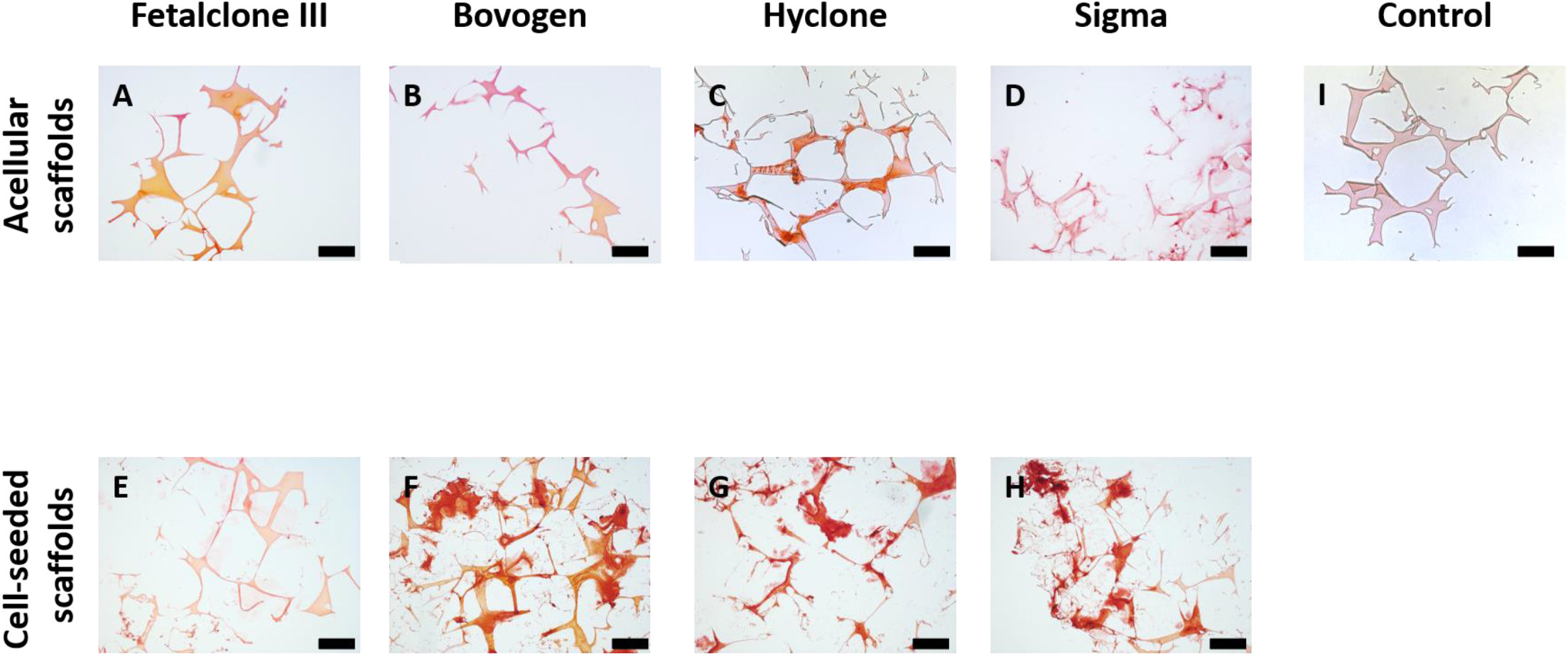
All constructs show mineral deposition with Alizarin Red staining. Acellular (A-D) and cell-seeded (E-H) 3D silk fibroin scaffolds after 4 weeks of culture in media containing different FBS types. In the medium containing Fetalclone III (A and E) and Bovogen (B and F) with low medium ALP activity, the mineral deposition happened exclusively in the presence of cells. In the Hyclone and Sigma containing medium groups with high medium ALP activity, substantial amounts of mineral deposition happened even if cells were not present (C, G, D, and H). Scale bar: 200 μm.

### 5 Enzymatic activity of ALP is declined in heat inactivated (HI) FBS

The ALP activity of HI FBS types and their corresponding control media containing 10% HI FBS was analysed. The heat inactivation process was able to decrease the ALP activity of FBS (Figure 6 A) compared to the non-HI FBS (Figure 1 A) by 88.05% (Fetalclone III), 94.77% (Bovogen), 95.61% (Hyclone), and 95.54% (Sigma), respectively. The ALP activity of 10% HI FBS in medium decreased further compared to concentrated FBS (Figure 6 B). To investigate the contribution of ALP activity of HI FBS in concentration of Pi in the medium, control media containing 10% HI FBS were supplemented with 10 mM β-GP. The concentration of Pi did not change in control medium supplemented with β-GP and 10% HI FBS after 48 hours of incubation compared to respective control medium (Figure 6 C). This result demonstrated that the ALP present in FBS can be deactivated through the HI process.

**Figure 6.**
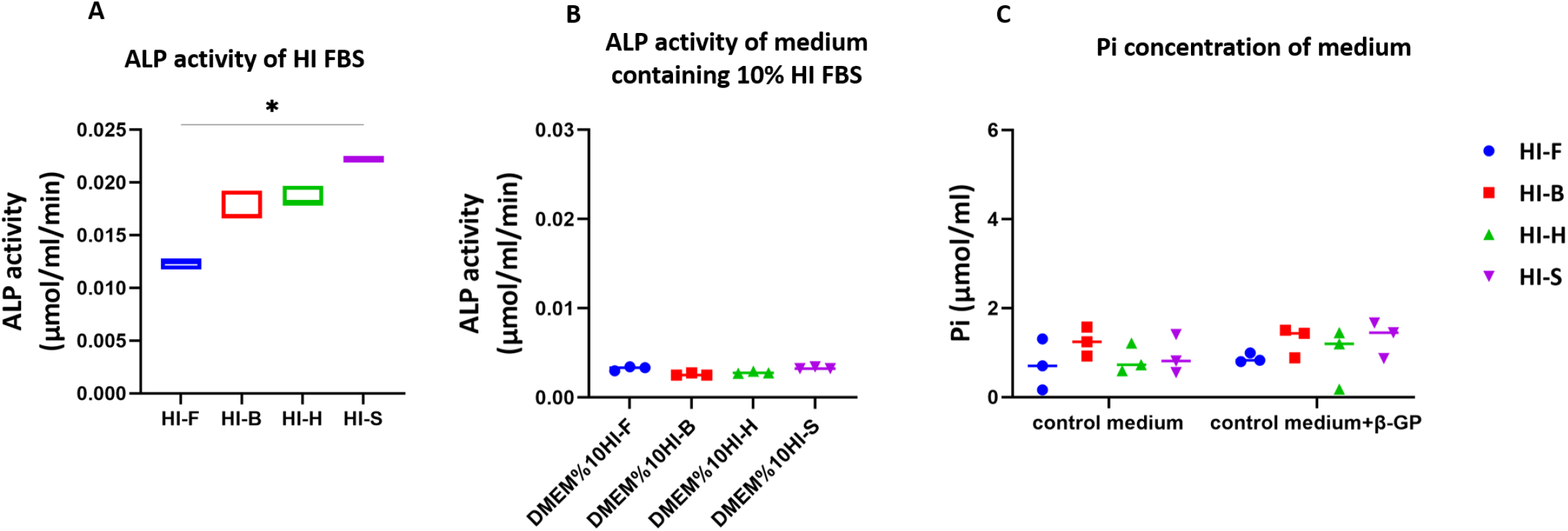
ALP activity of FBS decreased through HI process (A). A 10% dilution of FBS in media decreased ALP activity further and eliminated differences between the groups (B). 48 hours incubation of control medium supplemented with FBS and 10 mM β-GP resulted in no changes in Pi concentration in the medium which indicated the deactivation of ALP in HI FBS (C).

## Discussion

FBS was introduced more than 50 years ago as cell culture supplement for cellular growth as it contains crucial components for cell proliferation and maintenance including hormones, growth factors, vitamins, trace elements, and transport proteins [1,2,36]. However, the composition of FBS is not defined and consistent which could provoke significant differences in experimental outcomes and contribute to a low reproducibility of data [4,8,37]. Due to the disadvantages of using FBS in cell culture, it should be replaced by defined and more controlled media supplements. However, owing to time consuming and costly process of serum-free medium development, FBS is still a common cell culture supplement in cell culture practice. As such, researchers should at least be aware of a potential influence of FBS on their study outcomes, and, if needed, identify the influence of crucial factors.

It has previously been shown that FBS can affect the mineralization process in bone tissue engineering studies [7,8]. In the present study, ALP was investigated as a component present in FBS affecting in the mineralization process during *in vitro* bone-like tissue formation. We show that the inherent ALP activity of FBS could lead to significantly different conclusions about the osteogenic differentiation capability of cells and in particular about the amount of mineralized ECM deposition when performed with different FBS brands.

Bone tissue forms through two different procedures: endochondral ossification, which is a multistep process that requires the formation of cartilage template and its replacement with bone tissue, and intramembranous ossification through which bone tissue develops by the concentration of mesenchymal stromal cells (MSCs) that directly undergo osteogenic differentiation [38–40]. During intramembranous ossification, osteoblasts originating from MSCs deposit bone matrix through production of collagen type I fibrils and regulation of deposited minerals within the collagenous matrix [38]. To regulate the mineralization of collagenous matrix, osteoblasts express proteins including ALP which provides the phosphate required for mineralization process [12]. In bone tissue engineering and more precisely development *in vitro* bone models, the aim is to differentiate MSCs towards osteoblasts which produce the collagenous matrix and control matrix mineralization through the expression of non-collagenous proteins (NCPs) [30]. However, the presence of ALP - and possibly NCPs too - in FBS influences the whole osteogenic differentiation and mineralization process as we have shown here.

The presence of phosphatases in FBS was suggested in previous studies, as FBS showed the capability to hydrolyze β-GP and increase the phosphate concentration of medium in the absence of cells [41,42]. Among the proteins and phosphatases, ALP is a well-known one that is present in FBS and provides the cell culture media with free phosphate [29]. To the best of our knowledge, there is no evidence of the presence of other types of phosphatases that could hydrolyze β-GP. The four different brands of FBS tested in this study showed to differ in ALP activity in both concentrated and diluted state. Differences in the ALP activity of each FBS brand resulted in differences in the concentration of phosphate in the medium after 48 hours of incubation of FBS containing medium supplemented with β-GP; in the medium with low ALP activity (Fetalclone III), the lowest increased in phosphate concentration was detected. The amount of spontaneous mineralization depends on the ion concentration of the solution surrounding the substrate [43]. With the same basal medium being used, the initial calcium concentration of control medium was the same in all groups. As expected, differences in the ALP activity of FBS containing medium resulted in variation in Pi concentration in medium which further influences the calcium phosphate deposition.

The ALP activity of FBS containing medium also affected the cellular ALP activity. Though cells from the same vial (same donor, same passage) were used for the experiment, it seems that in the medium with high inherent ALP activity (Sigma), the cells showed lower ALP activity compared to the cells cultured in the medium with low ALP activity (Fetalclone III). This effect could be due to the calcium phosphate deposition because of the ALP activity of medium prior to expression of ALP from cells. In bone and calcifying cartilage, ALP is expressed early in the development, and is localized on the cell surface and on matrix vesicles. As the mineralized tissue matures, ALP expression and activity reduce [12,24]. Thus, the presence of calcium phosphate deposition prior to osteogenic differentiation of MSCs could have influenced the cellular ALP activity. Due to the complex and unknown composition of FBS, there is always a possibility of the presence of a component that influences the calcium phosphate deposition on cell-seeded/acellular scaffolds in addition to the cellular ALP activity. But a high inherent ALP activity of FBS could be a sign of how the cells might react or how the calcium phosphate deposits might develop in osteogenic differentiation cultures.

One possible way to avoid the influence of the ALP activity inherent to the various FBS brands is through the HI process. It is mostly used to destroy complement activity in serum through protein denaturation by heat and was also effective in reducing the ALP activity down to a base level that no longer was able to cleave substantial amounts of phosphate from β-GP. The HI process at the same time also influences the structural configuration of other heat-sensitive proteins which results in changes in their activity too [44]. These changes in turn can potentially affect the cellular behaviour including metabolic activity, proliferation and colony-forming units of hBMSCs [45]. An increase in cellular ALP activity has been reported when cells were cultured under HI serum supplemented medium [46]. This might be related to the reduction of the medium ALP activity but will need further investigation.

The present study was limited to four different FBS types and batch variation for each FBS types were not investigated here. But it can be expected that differences in ALP activity can be detected in other batches of the same FBS type and even other serum types, including human serum. Moreover, silk fibroin was the only biomaterial substrate tested in this study. The chemical structure of silk fibroin is similar to collagen type I which makes it an ideal environment for spontaneous mineralization, similarly as the collagen within the bone matrix. The ALP activity of FBS on mineralization process on other substrates might be different.

To avoid the effect of medium ALP activity and the other known/unknown components of FBS, development of defined and more controlled medium supplements is recommended. The information on existing defined media supplements is already available in databases, which could facilitate the process of formulating new media supplements [2]. So far, no general formula that suits all needs has been found. It seems as if these formulations are specific to the cell type, thus studying the factors impacting the specific cell behaviour is needed to develop such medium supplements.

## Conclusion

In this study, we have demonstrated that the ALP activity inherent to FBS influences both the cellular differentiation and the mineralization process, the two most important output parameters in bone tissue engineering. FBS types with differences in inherent ALP activity affected the calcium phosphate deposition in the presence and absence of cells. In media with high ALP activity, the amount of deposited calcium phosphate was higher compared to media with lower ALP activity. Moreover, the ALP activity of the medium affected the ALP activity of the cells; in media with higher ALP activity, the cellular ALP activity was reduced. Our results highlight the importance of considering the components present in FBS in tissue engineering studies. Generally, it is suggested that the development and optimization of specialized serum-free medium for tissue engineering applications should be advanced further.

## Conflict of interest

The authors declare that there is no conflict of interest.

## Acknowledgement

This work has been financially supported by the Dutch Ministry of Education, Culture and Science (Gravitation Program 024.003.013).

## Notes

### Competing Interest Statement

The authors have declared no competing interest.

